# Convolutional Neural Networks for Classifying Melanoma Images

**DOI:** 10.1101/2020.05.22.110973

**Authors:** Abhinav Sagar, Dheeba Jacob

## Abstract

In this work, we address the problem of skin cancer classification using convolutional neural networks. A lot of cancer cases early on are misdiagnosed leading to severe consequences including the death of patient. Also there are cases in which patients have other problems and doctors interpret it as skin cancer. This leads to unnecessary time and money spent for further diagnosis. In this work, we address both of the above problems using deep neural networks and transfer learning architecture. We have used publicly available ISIC databases for both training and testing our network. Our model achieves an accuracy of 0.935, precision 0.94, recall 0.77, F1 score 0.85 and ROC-AUC 0.861 which is better than the previous state of the art approaches.

## 1 Introduction

Skin cancer is the most widespread cancer diagnosed in the world. It is seen that if it can be diagnosed in its early phases, with choosing the appropriate treatment, survival rates are very good. Hence it is absolutely necessary to get to know at the earliest whether the symptoms of the patient correspond to cancer or not. Traditionally, doctors have been using their naked eye for skin cancer detection. However on many occasions this leads to not so accurate detection as people make mistakes. Even experts have a tough time saying it, especially when the cancer is at the very early stages. This is where computer vision can help in automating the whole pipeline.

A deep neural network can be trained on thousands of images of both the categories i.e. benign and malignant. By learning the non linear interactions, the model can tell whether a new image corresponds to a benign or malignant class. This automation could not only have a higher efficiency in avoiding both false positives and false negatives but also reduce time and manual spent on more productive work. Our model can be deployed on places which lack expert doctors. This work could dramatically change the healthcare and well being of humans in developing countries of Asia and Africa.

### 1.1 Image Classification

There are various image classification approaches which can be categorized under two heads - First one uses traditional machine learning algorithms like Decision Tree, Support Vector Machines, Fuzzy Measures etc and the other is based on deep learning like convolutional neural networks, autoencoders etc. Other than Artificial Neural Networks (ANNs), methods Decision Trees, Support Vector Machines, and Fuzzy Measures have been used.

Decision Trees is a machine learning method that works by calculating a class membership, which is done by partitioning a dataset into subsets repeatedly. This hierarchical classification permits approval and elimination of class labels at each stage. This method as a whole, has 3 major steps: Partition the nodes using the dataset, find all the terminal nodes, and allocate class labels to each terminal node. Support Vector Machines are used for a variety of machine learning applications. They work by building a hyper-plane or a set of hyper-planes, in a high-dimensional space. Good separation and classification is achieved when a hyper-plane has the largest distance to the nearest point of any class. Fuzzy measures another classification technique used. Multiple varied stochastic associations are determined in order to describe the image characteristics. The difference stochastic associations are then combined to form a set of properties.

### 1.2 Deep Learning

Deep learning is a set of machine learning methods that was inspired by information processing and distributed communication in networks of biological neurons. Deep learning predominantly involves development, training and utilization of artificial neural networks (ANNs). ANNs are networks of artificial neurons that are based on biological neurons. Every ANN has at least 3 layers: an input layer that takes the input, a hidden layer that trains on the dataset fed to the input layer, and an output layer that gives an output depending on the application.

Deep learning has been gaining wide acclaim because of the results it achieves that have never been seen in any other machine learning method. Convolutional Neural Networks (a type of ANNs), are extensively used for image-based applications, and have achieved better results than humans in object detection and classification.

### 1.3 Convolutional Neural Networks

CNNs are a kind of neural network which have proven to be very powerful in areas such as image recognition and classification. CNNs can identify faces, pedestrians, traffic signs and other objects better than humans and therefore are used in real time applications like robots and self-driving cars.

CNNs are a supervised learning method and are trained using labeled data given with the respective classes. CNNs learn the relationship between the input objects and the class labels and comprise two components: the hidden layers in which the features are extracted and, at the end of the processing, the fully connected layers that are used for the actual classification task. The hidden layers of CNN have a specific architecture consisting of convolutional layers, pooling layers and activation functions for switching the neurons either on or off. In a typical neural network, each layer is formed by a set of neurons and one neuron of a layer is connected to each neuron of the preceding layer while the architecture of hidden layers in CNN is slightly different. The neurons in a layer are not connected to all neurons of the preceding layer; rather, they are connected to only a small number of neurons from the previous layer. This restriction to local connections and additional pooling layers summarizing local neuron outputs into one value results in translation-invariant features. This results in a simpler training procedure due to less parameters and a lower model complexity.

## 2 Existing Work

There has been a lot of work published in the domain of skin cancer classification using deep learning and computer vision techniques. These works use a lot of different approaches including classification only, segmentation and detection, image processing using different types of filters etc.

(Esteva et al., 2017) separately used AdaBoost to classify skin lesions. (Xu et al., 2014) used different sets of features including type of lesion, texture, colour etc and neural networks for the making of a robust diagnosis system. The examples till now only showed algorithms using traditional machine learning techniques, but lately deep learning have proved to be more accurate. The reason is that it automates the feature extraction process completely. It is upto the algorithms to find the better features and train the model accordingly. In (Lopez et al., 2017) made a breakthrough on skin cancer classification by a pre-trained GoogleNet Inception v3 CNN model to classify 129,450 clinical skin cancer images including 3,374 dermatoscopic images. (Dorj et al., 2018) developed a convolutional neural network with over 50 layers on ISBI 2016 challenge dataset for the classification of malignant melanoma. In 2018, (Brinker et al., 2018) utilized a deep convolutional neural network to classify a binary class problem of dermoscopy images. (Rezvantalab et al., 2018) developed an algorithm using Support Vector Machines combined with a deep convolutional neural network approach for the classification of 4 diagnostic categories of clinical skin cancer images. (Codella et al., 2017) used a deep convolutional neural network to classify the clinical images of 12 skin diseases.

Our work builds on the aforementioned approaches. In this paper we have tackled skin cancer classification which is of binary type i.e. there are 2 classes present in the dataset benign and malignant. We have used the publicly available ISIC database for both training and testing the images. We have compared 5 backbone transfer learning architectures - VGG16, ResNet50, InceptionV3, MobileNet and DesneNet169. We have used familiar metrics for evaluating our results including confusion metric. Since there are equal numbers of images in both the classes, hence accuracy alone should be a good enough metric. Nevertheless we present other matric results including precision, recall, F1 score and ROC-AUC. Our work achieves an accuracy of 0.935, precision of 0.94, recall of 0.77, F1 score of 0.85 and ROC-AUC of 0.861 which is better than the previous state of the art approaches.

### 2.1 Skin Cancer Classification

A number of factors can make skin cancer recognition a challenging task. Some of these factors include flaws in image quality like uneven brightness, obstruction, and also the fact that there are many images which have similar shape, color and texture.

### 2.2 Dataset

We obtained a public dataset from ISIC website for skin cancer classification. We used the ISIC Archive Downloader to download the images. We used 3000 images for training and 600 images for validation of size 224 × 224. The images are distributed equally between training and validation sets which are shown below in Fig 2..

**Figure 1:**
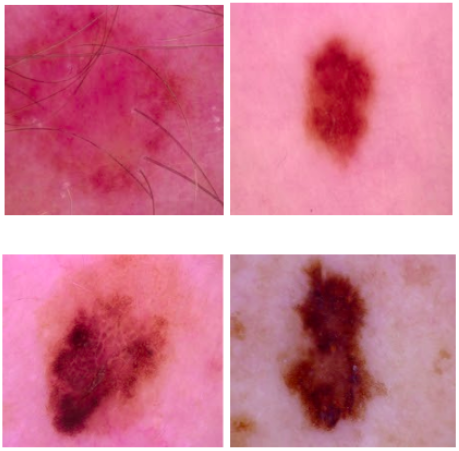
Benign vs malignant images

**Figure 2:**
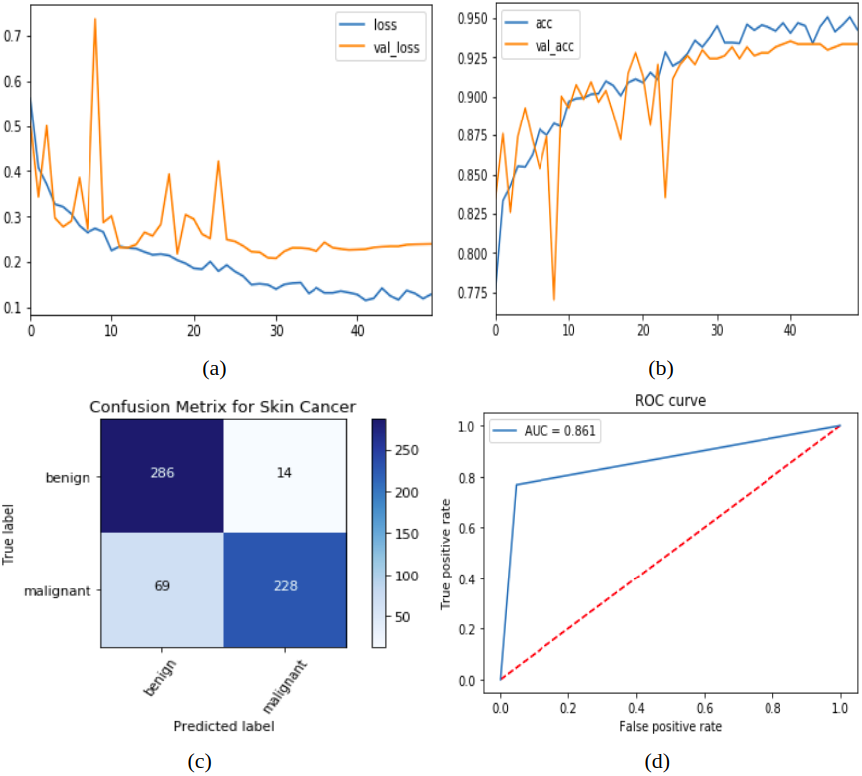
a) Loss vs epoch b) Accuracy vs epoch c) Confusion Matrix d) ROC-AUC curve

## 3 PROPOSED METHOD

We have used the concept of transfer learning for the classification. With transfer learning, instead of starting the learning process from scratch, the model starts from patterns that have been learned when solving a different problem. This way the model leverages previous learnings and avoids starting from scratch. In image classification, transfer learning is usually expressed through the use of pre-trained models. A pre-trained model is a model that was trained on a large benchmark dataset to solve a problem similar to the one that we want to solve. We used three pre-trained models-Inception v3, InceptionResNet v2 and ResNet 152 as the pre trained weights for our work.

### 3.1 Inception V3

Google’s Inception v3 architecture was re-trained on our dataset by fine-tuning across all layers and replacing top layers with one average pooling, two fully connected and finally the softmax layer allowing to classify 2 diagnostic categories. The size of input images was all resized to (224, 224) to be compatible with this model. Learning rate was set to 0.0001 and Adam was used for the optimizer.

### 3.2 InceptionResNet v2

InceptionResNet v2 architecture was re-trained on our dataset by fine-tuning across all layers and replacing top layers with one global average pooling, one fully connected and finally the softmax layer allowing to classify 2 diagnostic categories. The size of input images was all resized to (224, 224) to be compatible with this model. Learning rate was set to 0.0001 and Adam was used for the optimizer.

### 3.3 ResNet50

ResNet 50 architecture was re-trained on our dataset by fine-tuning across all layers and replacing top layers with one average pooling, one fully connected and finally the softmax layer allowing to classify 2 diagnostic categories. The size of input images was all resized to (224, 224) to be compatible with this model. Learning rate was set to 0.0001 and Adam was used for the optimizer. It uses identity mapping to map the inputs. This identity mapping does not have any parameters and is just there to add the output from the previous layer to the layer ahead. The identity mapping is multiplied by a linear projection to expand the channels of shortcut to match the residual. The Skip Connections between layers add the outputs from previous layers to the outputs of stacked layers. This results in the ability to train much deeper networks than what was previously possible.

### 3.4 MobileNet

The fundamental part of MobileNet is depthwise separable filters, named as Depthwise Separable Convolution. These convolutions layers which is a form of factorized convolutional factorize a standard convolution into a depthwise convolution called a pointwise convolution. In MobileNet, the depthwise convolution applies a single filter to each input channel. The pointwise convolution then applies the convolution operation to combine the outputs of the depthwise convolution.

### 3.5 DenseNet169

To solve the vanishing gradient problem, this architecture uses a simple connectivity pattern to ensure the maximum flow of information between layers both in forward and backward computation. The layers are connected in a way such that inputs from all preceding layers passes through its own feature-maps to all subsequent layers. To facilitate the down-sampling in the architecture, the entire architecture is divided into multiple densely connected blocks. The layers between these dense blocks are transition layers which perform convolution and pooling operations.

### 3.6 Transfer learning

Transfer learning is a popular method in computer vision that allows us to build accurate models faster. With transfer learning, instead of starting the learning process from scratch, we start from patterns that have been learned when solving a different problem. The advantages of using transfer learning are: Super simple to incorporate. Achieve the same or even better(depending on the dataset model performance quickly. There’s not as much labeled data required. Versatile uses cases from transfer learning, prediction, and feature extraction.

The proposed method can be summarized in the following seven points:

1. We split the dataset into two parts-training set and test set with 80 percent and 20 percent images respectively.
2. We used data augmentation like shearing, zooming, flipping and brightness change to increase the dataset size to almost double the original dataset size.
3. We tried with pre trained models like Inception v3, InceptionResNet v2, ResNet 50, MobileNet and DenseNet169 by fine tuning the last few layers of the network.
4. We used 50 percent dropout and batch normalization layers in between to reduce overfitting.
5. We used two dense layers with 64 neurons and 2 neurons respectively. The last layer is used for the classification with softmax as the activation function.
6. We used binary cross entropy as the loss function.
7. We trained the model for 20 epochs with a batch size of 32 by changing the hyper-parameters like learning rate, batch size, optimizer and pre-trained weights.

## 4 Experimental Results

In this section we present our findings. We plotted the loss vs epochs, accuracy vs epochs, confusion matrix for the classifier and ROC-AUC curve for the classifier. The plots are shown in Fig 4.

The details of experiment demonstrating the effect of training dataset size is shown in Table 1.

**Table 1:** Precision, recall F1 score values

| Train Size | Precision | Recall | F1 Score |
| --- | --- | --- | --- |
| 300 | 0.94 | 0.77 | 0.85 |
| 600 | 0.87 | 0.86 | 0.86 |

Next, we present our findings and show the validation of our trained models. In this paper, two types of major skin cancer categories are used. The evaluation and results of trained models is calculated by common classification metrics.

The ROC curve is calculated by plotting the sensitivity against 1-specificity and can be used to evaluate the classifier. The further the ROC curve deviates from the diagonal, the better the classifier.

We found that a batch size value of 64 gives better results when compared to that of 32 as shown in Table 2.

**Table 2:**
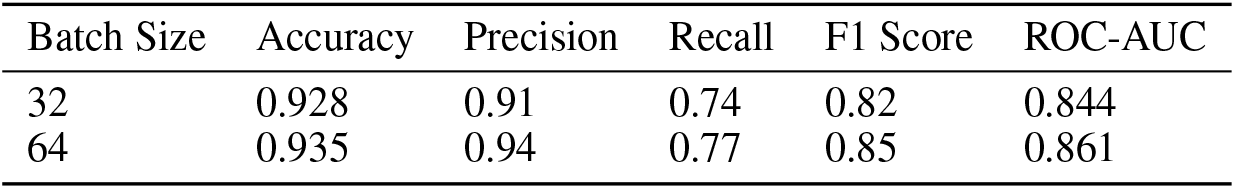
Effect of batch size on results

The effect of learning rate on results is shown in Table 3.

**Table 3:** Effect of learning rate on results

| Learning Rate | Accuracy | Precision | Recall | F1 Score | ROC-AUC |
| --- | --- | --- | --- | --- | --- |
| 0.0001 | 0.935 | 0.94 | 0.77 | 0.85 | 0.861 |
| 0.00001 | 0.915 | 0.88 | 0.72 | 0.79 | 0.813 |

We found that Adam as an optimizer performed better than Stochastic Gradient Descent (SGD) with momentum decay. The comparision of Adam vs SGD as optimizer is shown in Table 4.

**Table 4:** Comparison of Adam vs SGD as optimizer

| Optimizer | Accuracy | Precision | Recall | F1 Score | ROC-AUC |
| --- | --- | --- | --- | --- | --- |
| Adam | 0.935 | 0.94 | 0.77 | 0.85 | 0.861 |
| SGD | 0.903 | 0.92 | 0.71 | 0.80 | 0.830 |

We found that ResNet152 as the backbone gives better results when compared to InceptionV3, InceptionResNetV2, Mobilenet and Densenet169. The comparison of pre trained weights on results is shown in Table 5.

**Table 5:** Comparison of pre trained weights on results

| Pre Trained Model | Accuracy | Precision | Recall | F1 Score | ROC-AUC |
| --- | --- | --- | --- | --- | --- |
| ResNet50 | 0.935 | 0.94 | 0.77 | 0.85 | 0.861 |
| InceptionV3 | 0.893 | 0.92 | 0.71 | 0.80 | 0.839 |
| Inception ResNet v2 | 0.907 | 0.90 | 0.70 | 0.79 | 0.839 |
| MobileNet | 0.884 | 0.92 | 0.74 | 0.82 | 0.824 |
| Densenet169 | 0.903 | 0.93 | 0.73 | 0.82 | 0.847 |

The comparison with previous state of the art results is shown in Table 6.

**Table 6:** Comparison with previous work

| Work | Accuracy (percent) | ROC-AUC |
| --- | --- | --- |
| (Pomponiu et al., 2016) | 93.64 | - |
| (Codella et al., 2017) | 93.1 | - |
| (Haenssle et al., 2018) | - | 0.86 |
| (Han et al., 2018) | - | 0.83 |
| (Bi et al., 2017) | - | 0.915 |
| (Kawahara and Hamarneh, 2016) | 79.5 | - |
| (Lopez et al., 2017) | 81.33 | - |
| (Nasr-Esfahani et al., 2016) | 81 | - |
| Ours | 93.5 | 0.861 |

(Pomponiu et al., 2016) achieves a higher accuracy as they have used the complete dataset. Since our objective was to show comparable results with limited images, hence our accuracy is good enough considering the number of images on which our model is trained. (Bi et al., 2017) has much better ROC-AUC than ours as they have used an ensemble of CNNs for training, rather than ours which only used a single CNN architecture. Other than that our work archives state of the art results on skin cancer classification.

## 5 Conclusions

In conclusion, this study investigated the ability of deep convolutional neural networks in the classification of benign vs malignant skin cancer. Our results show that state-of-the-art deep learning architectures trained on dermoscopy images (3600 in total composed of 3000 training and 600 validation) outperforms dermatologists. We showed that with use of very deep convolutional neural networks using transfer learning and fine-tuning them on dermoscopy images, better diagnostic accuracy can be achieved compared to expert physicians and clinicians. Although no preprocessing step is applied in this paper, the experimental results are very promising. These models can be easily implemented in dermoscopy systems or even on smartphones in order to assist dermatologists. More diverse datasets (varied categories, different ages) with much more dermoscopy images and balanced samples per class is needed for further improvement. Also using the metadata of each image can be useful to increase the accuracy of the model.

## Acknowledgments

We would like to thank Nvidia for providing the GPUs.

